# Accelerating Protein-Protein Interaction screens with reduced AlphaFold-Multimer sampling

**DOI:** 10.1101/2024.06.07.597882

**Authors:** G. Bellinzona, D. Sassera, A.M.J.J Bonvin

## Abstract

**Motivation:** Discovering new protein-protein interactions (PPIs) across entire proteomes offers vast potential for understanding novel protein functions and elucidate system properties within or between an organism. While recent advances in computational structural biology, particularly AlphaFold-Multimer, have facilitated this task, scaling for large-scale screenings remains a challenge, requiring significant computational resources.

**Results:** We evaluated the impact of reducing the number of models generated by AlphaFold-Multimer from five to one on the method’s ability to distinguish true PPIs from false ones. Our evaluation was conducted on a dataset containing both intra- and inter-species PPIs, which included proteins from bacterial and eukaryotic sources. We demonstrate that reducing the sampling does not compromise the accuracy of the method, offering a 5 time faster, efficient, and environmentally friendly solution for PPI predictions.

**Availability:** The code used in this article is available at https://github.com/MIDIfactory/AlphaFastPPi. Note that the same can be achieved using the latest version of AlphaPulldown available at https://github.com/KosinskiLab/AlphaPulldown.

## Introduction

Mapping protein-protein interactions (PPIs) at the whole proteome level holds significant promise for uncovering new protein functions and elucidating local and global system properties within biological systems. Within an organism, PPIs are essential for vital cellular processes, such as signal transduction, metabolic pathways, and gene regulation. Interactions between different organisms, such as host-pathogen interactions, are also mediated by PPIs and are critical for understanding broader biological mechanisms.

Traditionally, studying these interactions has been laborious and often limited by the constraints of experimental techniques. The advancement of computational tools for predicting protein structures, particularly the recent introductions of Alphafold2 (Jumper *et al*., 2021) and AlphaFold-Multimer (Evans et al., 2022), has significantly enhanced our ability to study PPIs.

AlphaFold-Multimer leverages the power of GPUs to model PPIs, making the prediction process not only accurate but also relatively fast. Since its release, several optimized pipelines have been introduced to accelerate the computational workflow even further. These improvements include the separation of CPU and GPU stages to optimize processing efficiency and providing faster multisequence alignment (MSA) search options (Mirdita *et al*., 2022). Additionally, a number of tools have been developed to help with specific AlphaFold-Multimer (Evans et al., 2022) applications, including AlphaPulldown (Yu *et al*., 2022), a python package which streamlines the screening of PPIs. Despite these advancements, large-scale PPI screening remains challenging due to the substantial requirements for computing time, disk space, and GPU resources. These constraints can limit the accessibility and scalability of AlphaFold-Multimer and AlphaPulldown (Yu *et al*., 2022) for large scale studies.

To help overcome this issue, we evaluated the impact of reducing AlphaFold-Multimer sampling from the default settings of five models to a single one on its ability to recognise true PPIs. To achieve this, we developed an AlphaPulldown v.1.0 -based python wrapper called AlphaFastPPI. This change aims to address several critical needs: (i) it enhances the accessibility for high-throughput PPI screenings for users with limited computational capabilities; (ii) it conserves computational resources by lowering the time required by a factor 5, which is crucial to enable more sustainable large-scale analyses; and (iii) importantly, it should do so without compromising the accuracy required for confident predictions.

We tested this reduced sampling on a dataset consisting of intra- and inter-species PPIs comprising bacterial and eukaryotic proteins. This work extends the accessibility of AlphaFold-Multimer’s predictive capabilities while reducing environmental impact and resource wastage, making it a more inclusive and sustainable method.

## Code implementation

First, Multiple Sequence Alignments (MSA) and structural template features are generated using the AlphaPulldown v.1.0.4 (Yu *et al*., 2022) script create_individual_features.py on CPUs.

An *ad-hoc* pipeline, AlphaFastPPI, was developed at the time because AlphaPulldown v.1.0.0 did not support single model predictions. AlphaFastPPi, which generates a single model for each PPI on GPUs when available, otherwise CPUs are used. Since it has been demonstrated that MSA pairing does not improve AlphaFold performance, only unpaired MSAs are used to increase speed (Yin *et al*., 2022). AlphaFastPPi offers two modalities, similar to AlphaPulldown v.1.0.4 (script run_multimer_jobs.py) (Yu *et al*., 2022): pull-down and all-versus-all. In pull-down mode, users provide a list of one or more proteins as ‘baits’ and another protein as ‘candidates’, which are potential interaction partners. Then, AlphaFold-Multimer evaluates each bait protein against each candidate protein, generating a single model for each pair. In all-versus-all mode, the software computes all possible combinations among the proteins in the provided list, generating a single model for each pair. After modeling, model quality scores such as pDockQ (Bryant *et al*., 2022), ipTM, ipTM+pTM, and average plDDT are calculated for each predicted PPI and the results are stored in a table for further analysis. The workflow is shown in Figure 1.

**Figure 1.**
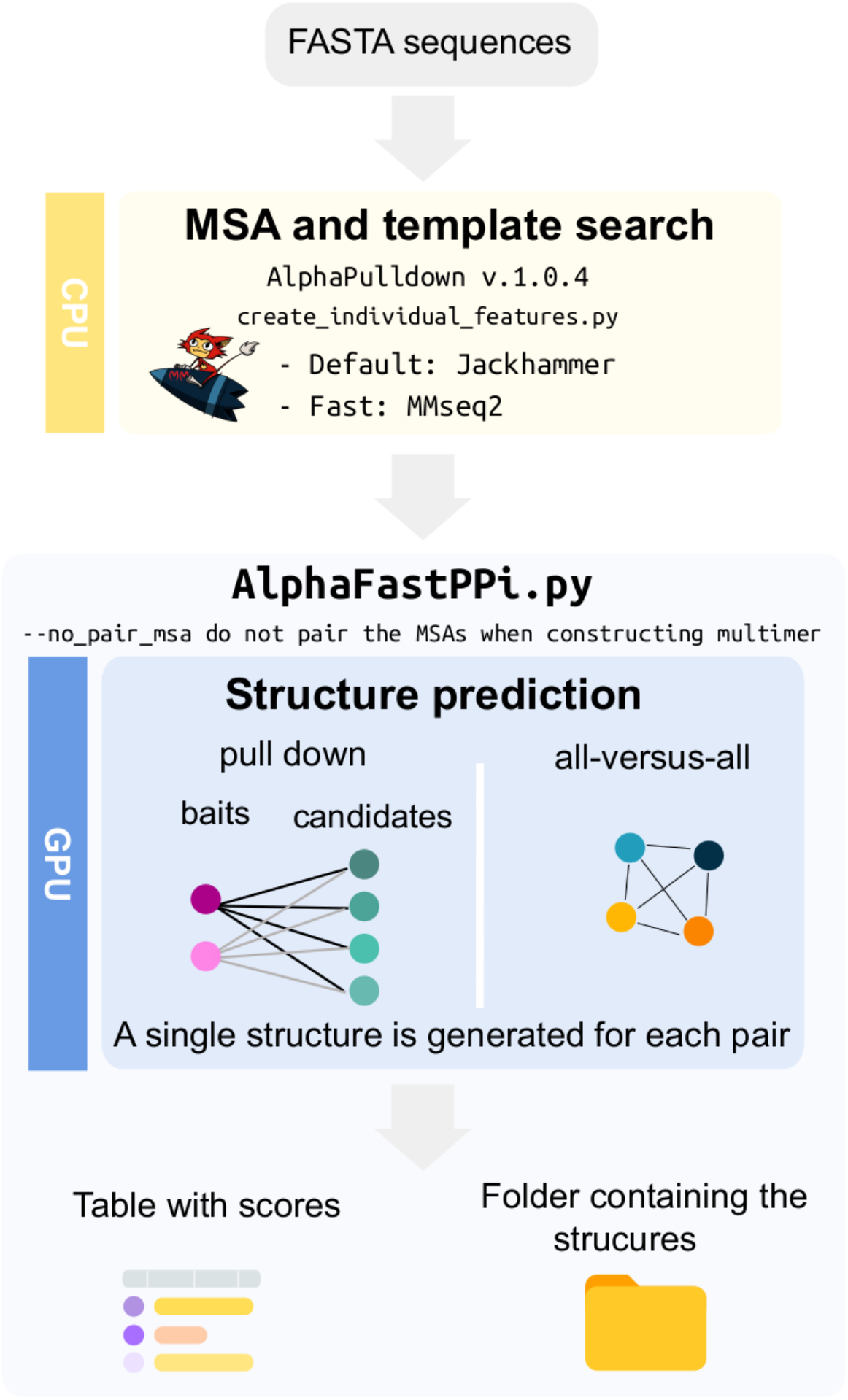
Overview of the AlphaFastPPi workflow.

Note that the same result can now be achieved by using `run_multimer_job.py` from AlphaPulldown v.1.0.4 using options `--num_predictions_per_model=1, -- model_names=model_1_multimer_v3, --num_cycle=1, --nopair_msa`.

## Benchmark dataset

A test set was built by combining both experimentally validated positive PPIs and a set of negative interactions taken from the Negatome database (Blohm *et al*., 2014) (Supplementary Table 1). The positive test set was composed of 286 PPIs from *Mycoplasma pneumoniae* (Elfmann et al., 2022) and 142 heterodimeric protein complexes between *Arabidopsis thaliana* and *Serendipita indica* (Osborne *et al*., 2023). The negative data set consisted of an equal number (428) of true negatives composed by randomly selecting non-interacting protein pairs from the Negatome (Blohm *et al*., 2014).

## Results

We evaluated the impact of reducing the sampling from the default of five models to a single model on the method’s performance in distinguishing interacting from non-interacting protein pairs. This comparison was conducted using experimentally-proven interacting and non-interacting protein pairs. All the predictions were performed on NVIDIA A100 SXM6 64GB GPUs. While GPU acceleration significantly speeds up the code, it is not a strict requirement. If no GPU is detected, the tool will automatically turn down to use CPU resources at the cost of increased computational time. As expected, reduced sampling allowed us to save approximately 4.7 times the GPU hours, translating to over 8 days in total for screening 856 interactions, and approximately 5.0 times the disk space, saving more than 2 TB of disk space. Although various scoring metrics (e.g. ipTM, pDockQ), have been developed to evaluate the quality of a model, none have been explicitly designed to address this specific question of whether two proteins are interacting or not. However, among these metrics, pDockQ displayed some capability in addressing this task (Bryant *et al*., 2022). When considering default sampling (five models), only the model with the highest pDockQ among the five generated was considered to compare the distribution of values with those from collected predicting just one model for each PPI. The same approach was applied for ipTM. We did not observe any significant difference between the two methods considering both scores. This suggests that lowering the number of predictions from five to a single one does not affect the discriminatory power in distinguishing between interacting and non-interacting pairs (Figure 2).

**Figure 2.**
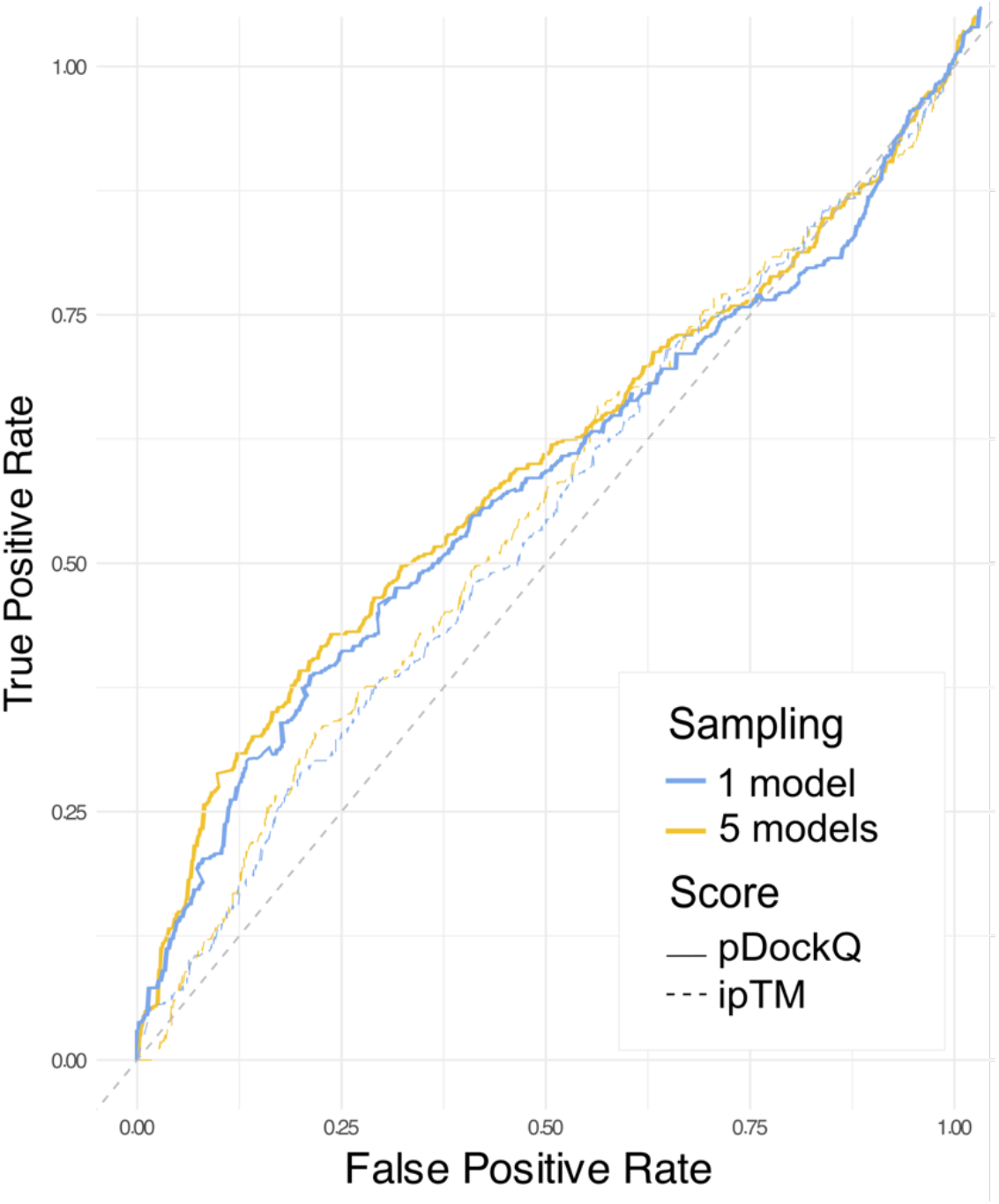
ROC curves comparing 5 models (yellow) and 1 model (blue) based on pDockQ and ipTM. The ROC curves using both the pDockQ (solid line) and ipTM (dashed line) scores to distinguish true from false interactions are shown. DeLong’s test was performed to compare 5 models and 1 model sampling for both ipTM and pDockQ, indicating no statistically significant difference between the AUCs (pDockQ Z=0.53849, p-value=0.428; ipTM: Z=0.41, p-value=0.328).

Consistent with previous findings (Bryant *et al*., 2022), pDockQ performed better than ipTM. When applying a pDockQ cut-off of 0.23, deemed sufficient to ensure a reasonable model quality (Burke *et al*., 2023), both AlphaPulldown and AlphaFastPPi achieved a specificity above 70%, with sensitivity around 24% on our benchmark dataset. Increasing the pDockQ threshold to 0.5 resulted in a specificity above 90%, although the sensitivity dropped below 10%. Moreover, interspecies interactions (i.e between *A. thaliana* and *S. indica* proteins in this benchmark) showed lower confidence scores compared to intra-species interactions (i.e among *M. pneumoniae* proteins). It should be mentioned that massive sampling, which has been demonstrated to effectively increase the quality of models (Wallner, 2023), will likely also enhance the method’s ability to distinguish between interacting and non-interacting protein pairs. However, this approach is not applicable for large-scale studies due to its substantial computational requirements.

## Conclusions

We showed that reducing the sampling of AlphaFold-Multimer from the default value of five models to one, reduces computational resources consumption by a factor 5 while maintaining a competitive performance in predicting interacting pairs. This aligns with the needs of the scientific community, ranging from conducting quick preliminary analyses, offering a way for a rapid and effective means of narrowing down potential interactions for further investigation, to those involved in extensive studies with limited resources. Additionally, it supports extensive studies with limited resources and promotes more sustainable computational research on PPIs.

## Availability

The code used in this article is available on GitHub https://github.com/MIDIfactory/AlphaFastPPi

